# Bouncer’s receptor on sperm: Investigating sperm-egg compatibility in fish

**DOI:** 10.1101/2024.12.13.628393

**Authors:** Andreas Blaha, Andrea Pauli

## Abstract

Fertilization requires the successful binding and fusion of sperm and egg. In zebrafish, sperm-egg binding is mediated by the Spaca6-Izumo1-Tmem81 complex on sperm interacting with Bouncer on the egg. We previously found that expressing medaka Bouncer on zebrafish eggs and *vice versa* allows hybridization between medaka and zebrafish, two species that can normally not hybridize. Here, we tested whether providing zebrafish Spaca6 and Izumo1 on medaka sperm and vice versa enables cross-species compatibility from the side of the sperm. To this end, we generated *spaca6* and *izumo1* knock-out (KO) lines in medaka, which are male sterile, and introduced zebrafish *spaca6* and *izumo1* transgenes. Transgenic medaka males did not fertilize zebrafish or medaka eggs with zebrafish Bouncer. Similarly, zebrafish males expressing medaka Spaca6 and Izumo1 failed to fertilize zebrafish eggs presenting medaka Bouncer. Unexpectedly, providing either full-length medaka Spaca6 or the Bouncer binding site of medaka Izumo1 in zebrafish sperm rescued the sterility of *spaca6* and *izumo1* KO, respectively. Therefore, medaka Spaca6 and Izumo1 can interact with zebrafish Bouncer when paired with their zebrafish sperm complex members underscoring the nuanced interplay between molecular restrictions and compatibilities during sperm-egg interaction across teleosts.

## Introduction

Fertilization is the final step in sexual reproduction and marks the beginning of a new individual. Despite its fundamental importance, the molecular underpinnings governing sperm-egg binding and fusion in vertebrates are still poorly understood. One particular question that has remained unanswered is how incompatibilities between sperm and egg are encoded at the molecular level.

Decades of meticulous genetic experiments and recent advances in artificial intelligence have provided important insights into the mechanisms of sperm-egg interaction in vertebrates (Lu and Ikawa, 2022; Deneke et al., 2024; Elofsson et al., 2024). To date, three egg proteins are known to be essential for vertebrate fertilization. CD9 and JUNO are required on mammalian eggs (Miyado et al., 2000; Kaji et al., 2000; Le Naour et al., 2000; Rubinstein et al., 2006; Bianchi et al., 2014). JUNO is a mammalian-specific folate receptor homolog and directly interacts with mammalian IZUMO1 on the sperm (Bianchi et al., 2014). Fish lack JUNO but express the unrelated Bouncer protein on the egg, which is required for sperm-egg binding (Herberg et al., 2018; Gert et al., 2023; Yoshida et al., 2024). Bouncer is a small GPI-anchored Ly6/uPAR protein with a characteristic three-finger fold (Herberg et al., 2018). Strikingly, expression of medaka (*Oryzias latipes*) Bouncer on zebrafish (*Danio rerio*) eggs and *vice versa* is sufficient to allow hybridization between medaka and zebrafish, two species whose gametes cannot cross-fertilize even *in vitro* (Herberg et al., 2018; Gert et al., 2021). Detailed analysis of Bouncer’s surface residues and glycosylation pattern identified molecular signatures that delineate Bouncer recognition by either medaka or zebrafish sperm (Gert et al., 2023). However, both zebrafish and medaka sperm are compatible with seahorse and fugu Bouncer despite the considerable sequence divergence and evolutionary distance (Gert et al., 2023). This suggests that Bouncer’s primary function is to enable binding and subsequent fusion between compatible sperm and egg.

Recently, a study guided by artificial intelligence identified Bouncer’s receptor on sperm, a conserved trimeric complex formed by Spaca6, Izumo1 and the newly characterized factor Tmem81 (Deneke et al., 2024). Bouncer was shown to bind in a cleft formed by Spaca6 and Izumo1 thereby bridging sperm and egg (**Fig. 1A**; Deneke et al., 2024). Therefore, the seemingly parallel systems for sperm-egg adhesion in fish and mammals converged since Bouncer and JUNO bind the same orthologous sperm complex (Deneke et al., 2024; Elofsson et al., 2024).

**Figure 1.**
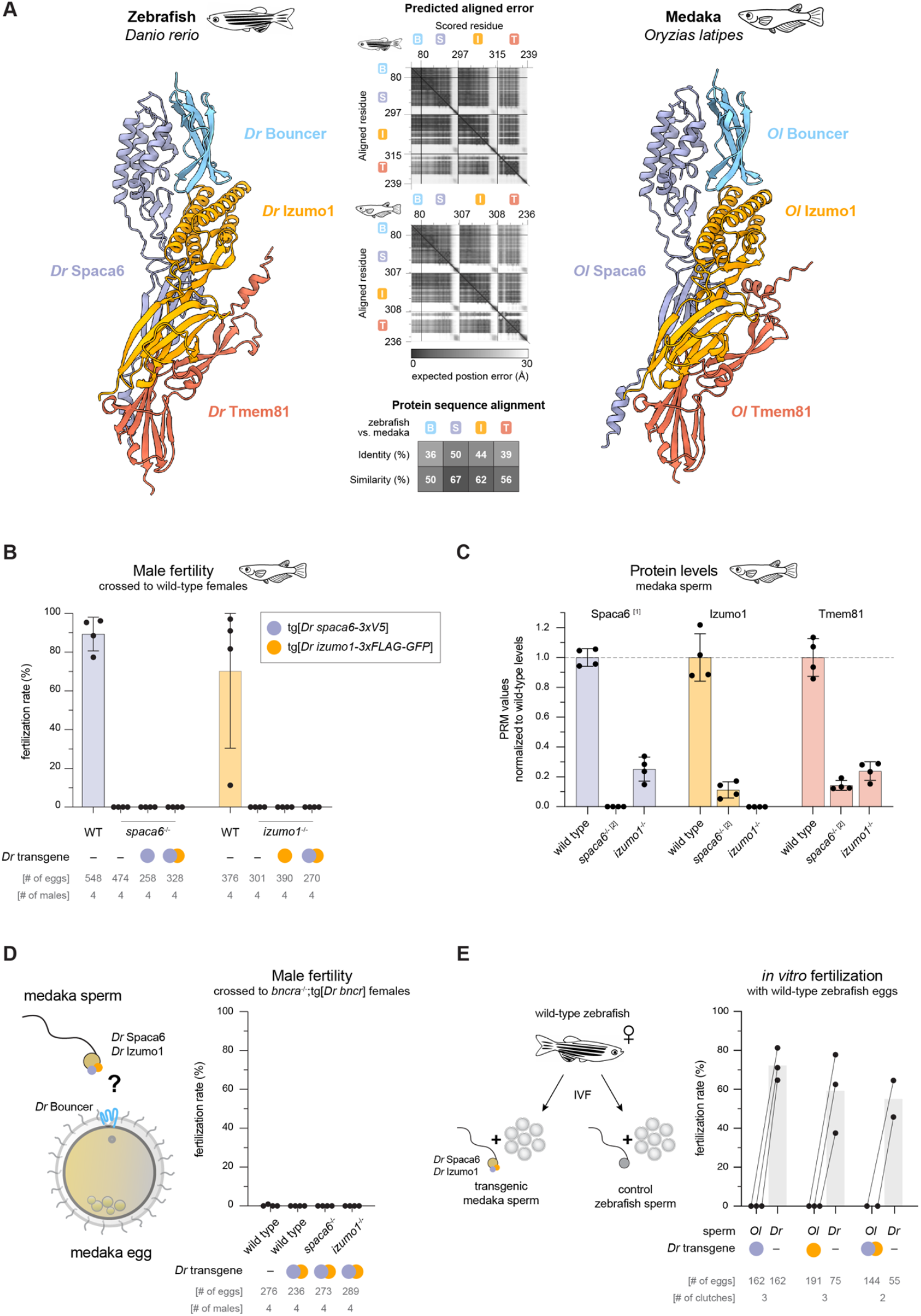
Medaka Spaca6 and Izumo1 are essential for male fertility and cannot be substituted by zebrafish *spaca6* and *izumo1* transgenes. **A** AlphaFold2-Multimer predicts a tetrameric complex of Spaca6, Izumo1, Tmem81 and Bouncer both in zebrafish (*Danio rerio, Dr*) (left) and in medaka (*Oryzias latipes, Ol*) (right). Cartoon models of the ectodomains as predicted in the relaxed top-scoring model. Predictive confidence as visualized by the predicted aligned error (PAE) plot (center top). Zebrafish model and PAE plot modified from Deneke et al., 2024. The calculated amino acid identity and similarity scores between zebrafish and medaka proteins from Needleman-Wunch alignments are depicted in shades of grey (center bottom). **B** *In vivo* fertility of mutant medaka males compared to their wild-type siblings was assessed by co-housing individual males with wild-type females and determining the fertilization rate of all eggs of that cross. Transgenic expression was driven by the upstream sequence of testis-expressed *Ol dcst2*. **C** Targeted mass spectrometry via parallel reaction monitoring (PRM) of Spaca6, Izumo1 and Tmem81 protein in sperm lysate from wild-type, *spaca6*-/- or *izumo1*^-/-^ males. To correct for sample input, six control proteins deemed constant were quantified as well and used to scale the values of the Spaca6, Izumo1 and Tmem81 peptides. The plotted values are the normalized peptide counts relative to the wild-type mean. **D** Assessment of male fertility as in (B) but with medaka females expressing solely zebrafish Bouncer protein (*bncra*^-/-^; tg[*Dr bncr*]). **E** *In vitro* fertilization (IVF) of wild-type zebrafish eggs with medaka sperm expressing zebrafish *spaca6* and/or *izumo1*. The same clutch was split and tested with wild-type zebrafish sperm to account for clutch-to-clutch variation.**[1]** One of the four Spaca6 peptides measured was excluded from the final quantification. Refer to **Fig. S1C** for the considerations leading to the exclusion. **[2]** Two of four *spaca6*^-/-^ samples were collected from *spaca6*^-/-^; tg[*Dr spaca6*] males and one sample from *spaca6*^-/-^; tg[*Dr spaca6*]; tg[*Dr izumo1*] males.

The identification of Bouncer’s receptor on sperm gave us the opportunity to ask whether Bouncer’s capacity to encode sperm-egg compatibility is mirrored by its receptor on sperm, Izumo1 and Spaca6. Can species compatibility of zebrafish or medaka sperm be switched by expressing the orthologous Izumo1 and/or Spaca6? We introduced zebrafish *spaca6* and *izumo1* in medaka sperm and *vice versa* and tested their compatibility with the heterospecific Bouncer on medaka and zebrafish eggs. Overall, our work provides new insights into how sperm-egg compatibilities are encoded on a molecular level.

## Results

### The conserved fertilization complex is predicted to form in medaka

Recently, we discovered a sperm complex consisting of the fertility factors Spaca6, Izumo1 and Tmem81, which is conserved across vertebrates and acts as Bouncer receptor in zebrafish (Deneke et al., 2024) (**Fig. 1A**). Because all four factors have direct medaka orthologs, we hypothesized that this complex also forms in medaka. Indeed, AlphaFold2-Multimer predicted with high confidence a tetrameric complex of medaka Spaca6, Izumo1, Tmem81 and Bouncer (Evans et al., 2022; Abramson et al., 2024). This complex adopts the same overall fold and conformation as the zebrafish complex despite the relatively high divergence in sequence (**Fig. 1A**). Given that exchanging zebrafish Bouncer for its medaka homolog is sufficient to allow medaka sperm to fertilize a zebrafish egg and *vice versa* (Herberg et al., 2018; Gert et al., 2021, 2023), we asked whether substituting Bouncer’s direct interaction partners on sperm, Spaca6 and Izumo1, with their orthologs is sufficient for the sperm to recognize heterospecific Bouncer.

### Izumo1 and Spaca6 are essential for male fertility in medaka

In order to replace medaka Izumo1 and Spaca6 for their zebrafish ortholog, we generated medaka *spaca6* and *izumo1* knock-out (KO) lines by CRISPR/Cas9-mediated mutagenesis (**Fig. S1A** and **B**). Consistent with their known role in male fertility, both *spaca6*^-/-^ and *izumo1*^-/-^ males were sterile solidifying that Spaca6 and Izumo1 are core fertilization factors across vertebrates (**Fig. 1B**).

We previously found that in zebrafish, the stability of all three proteins of the sperm complex, Spaca6, Izumo1 and Tmem81, is interdependent in mature sperm: If one factor is ablated, the other two will be lost as well (Deneke et al., 2024). In contrast, the stability of murine IZUMO1 is not dependent on any of the known sperm fertilization factors (Noda et al., 2020; Barbaux et al., 2020; Inoue et al., 2021). To ask whether the medaka factors behave like the zebrafish or murine orthologs, we adapted the targeted mass spectrometry method (parallel reaction monitoring (PRM)) developed for zebrafish sperm (Deneke et al., 2024) to medaka sperm to quantify the relative protein levels of medaka Spaca6, Izumo1 and Tmem81. This modified set-up could confidently detect medaka Spaca6, Izumo1 and Tmem81 in mature wild-type sperm, and as expected, Spaca6 and Izumo1 were no longer detected in the respective mutant (**Fig. 1C**). In addition, knocking-out *spaca6* or *izumo1* also caused a marked decrease of the other two complex members (**Fig. 1C**). While this observation resembles the situation in zebrafish (Deneke et al., 2024), the level of the remaining factors was higher in medaka (∼15-30% of wild-type level), whereas the co-regulated factors are almost completely lost in zebrafish (undetectable or <10%). This suggests that interdependence of Spaca6-Izumo1-Tmem81 is broadly conserved in teleosts albeit less pronounced in medaka.

Taken together, these results show that medaka Spaca6 and Izumo1 are required for male fertility and that their stability is interdependent suggesting that the Spaca6-Izumo1-Tmem81 complex also forms in medaka.

### Medaka Spaca6 and Izumo1 cannot be substituted by their orthologous zebrafish transgenes

To replace medaka *spaca6* and *izumo1* with their zebrafish orthologs, we introduced transgenes encoding tagged versions of zebrafish Spaca6 and Izumo1 into the respective medaka mutant line. Additionally, we generated lines that expressed both zebrafish factors in either mutant background. Medaka males carrying zebrafish *spaca6* or *izumo1* transgenes did not fertilize wild-type medaka eggs, even when providing both zebrafish factors as transgenes (**Fig. 1B**). This was expected given that zebrafish sperm can only inefficiently fertilize medaka eggs *in vitro* due to the incompatible Bouncer on the egg (Gert et al., 2021). Hence, we asked whether the transgenic medaka sperm can fertilize medaka eggs presenting zebrafish Bouncer. These eggs - once exogenously activated with calcium ionophore - are effectively fertilized by zebrafish sperm *in vitro* (Gert et al., 2021). Therefore, we assessed the fertility of the transgenic males when crossed to *bncra*^-/-^; tg[*Dr bncr*] females (**Fig. 1D**). Neither *spaca6*^-/-^ nor *izumo1*^-/-^ males carrying zebrafish *spaca6* and/or *izumo1* transgenes were able to fertilize the zebrafish Bouncer-expressing medaka eggs. This negative result could be due to the observed interdependence of the Spaca6-Izumo1-Tmem81 complex, since zebrafish Spaca6 and Izumo1 could be incompatible with medaka Tmem81 and destabilized. Therefore, we tested transgenic medaka sperm in the wild-type background since in this scenario the interaction of zebrafish Spaca6-Izumo1-Bouncer could provide adhesion across the gametes alongside the endogenous medaka sperm complex. However, also these males did not fertilize medaka eggs presenting zebrafish Bouncer (**Fig. 1D**).

Finally, we hypothesized that the transgenic medaka sperm can fertilize wild-type zebrafish eggs, since expressing medaka Bouncer is sufficient to enable wild-type medaka sperm to fertilize zebrafish eggs (Herberg et al., 2018; Gert et al., 2023). To this end, we performed IVF with zebrafish eggs and the transgenic medaka sperm. However, none of the tested sperm was able to fertilize wild-type zebrafish eggs even though the IVF performed on the same clutches with zebrafish sperm was successful (**Fig. 1E**).

These results can have several explanations: (1) The zebrafish factors are not or only lowly present as proteins in sperm. The medaka *dcst2* promoter might be driving insufficient expression of the transgenes, which would be in contrast to the previously verified expression by the equivalent zebrafish *dcst2* promoter sequence (Deneke et al., 2024). Alternatively, the proteins may not be present in mature sperm due to the observed interdependence. (2) Even if the zebrafish proteins were present in medaka sperm, they may require and lack compatibility with endogenous medaka sperm factors to function – first and foremost with medaka Tmem81. (3) If zebrafish Spaca6 and Izumo1 were present on the surface of medaka sperm at sufficient levels, these results would suggest that simply enabling Bouncer binding from the side of the sperm was not sufficient to proceed with gamete fusion.

Despite these limitations, the presented experiments show that introducing transgenes encoding zebrafish Spaca6 and Izumo1 is not sufficient to allow medaka sperm to fertilize zebrafish Bouncer-presenting eggs.

### Medaka Spaca6 can rescue zebrafish *spaca6* mutants whereas medaka Izumo1 cannot replace zebrafish Izumo1

Since the Spaca6 and Izumo1 substitutions in medaka remained ambiguous, we tested the reverse scenario. To this end, we introduced medaka *spaca6* and *izumo1* transgenes into zebrafish *spaca6* and *izumo1* mutant backgrounds. To our surprise, transgenic medaka Spaca6 fully rescued the sterility of *spaca6* KO zebrafish (**Fig. 2A**). This result shows that medaka Spaca6 is present in the mature sperm. It also suggests that medaka Spaca6 is integrated into the zebrafish complex and stabilizes endogenous zebrafish Izumo1 and Tmem81, and that it binds zebrafish Bouncer when forming the composite binding site together zebrafish Izumo1.

**Figure 2.**
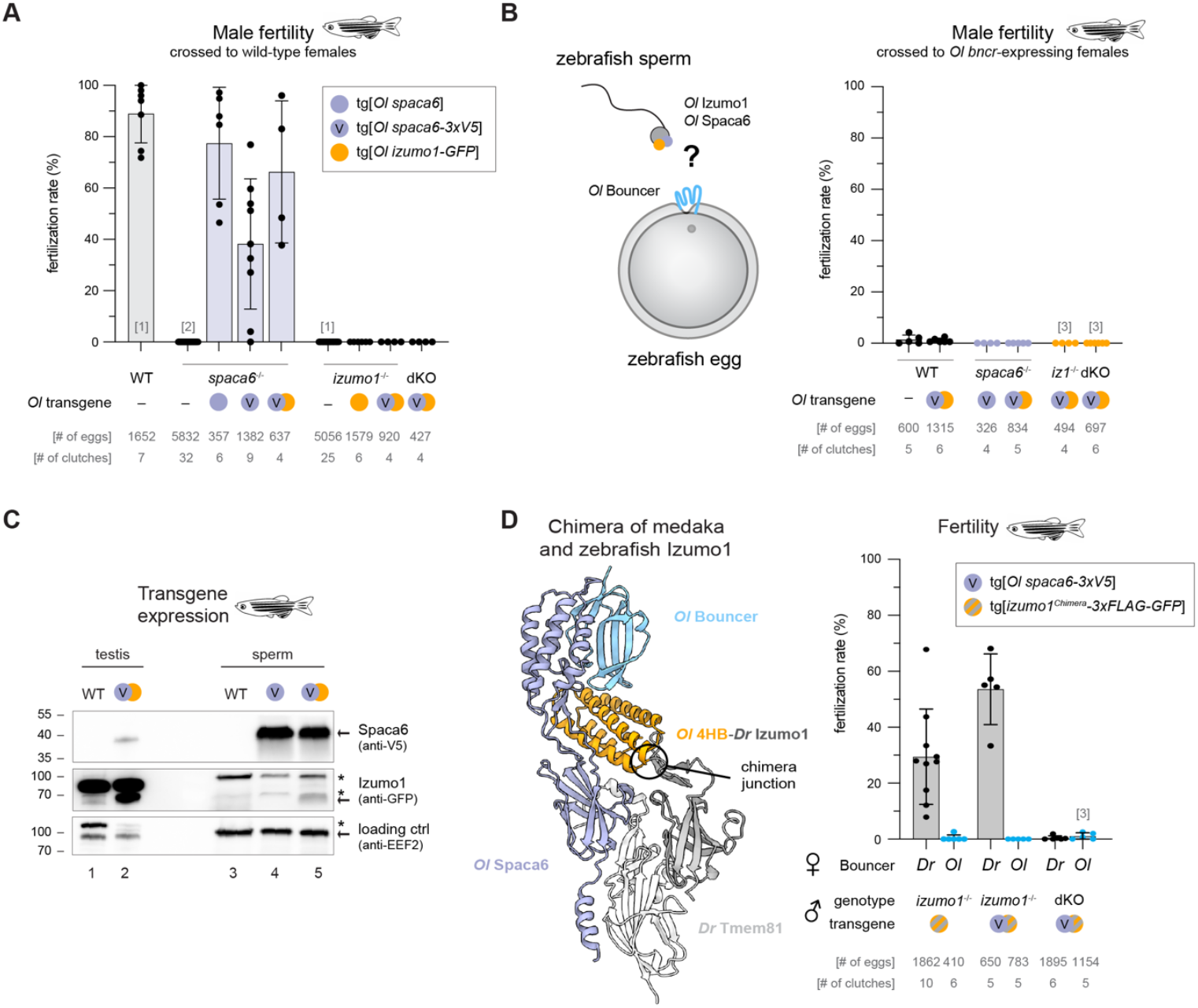
Providing the medaka Bouncer binding site of either Spaca6 or Izumo1 is compatible with zebrafish Bouncer but not medaka Bouncer. **A** To assess whether medaka Spaca6 or Izumo1 can rescue zebrafish *spaca6* or *izumo1* KO or double KO (dKO), transgenic males of the indicated genotypes were crossed to wild-type zebrafish females. The fraction of developing embryos per clutch (fertilization rate) is plotted compared to wild-type and mutant males. **B** *In vivo* fertility rates were scored as in (A) but with zebrafish *bncr*^-/-^; tg[*GFP-bncr*(*Ol* Top)] females (generated in Gert et al., 2023) to test the compatibility of the transgenic sperm with medaka Bouncer. **C** Western blot analysis of transgenic testis and sperm lysate expressing medaka Spaca6-3xV5 and Izumo1-GFP. Genotype per lane: (1 and 3) wild-type, (2) *spaca6*^-/-^; tg[*Ol spaca6-3xV5*]; tg[*Ol izumo1-GFP*]; tg[*Ol Tmem81-ALFA*], (4) *spaca6*^-/-^; tg[*Ol spaca6-3xV5*], (5) *spaca6*^-/-^; *izumo1*^-/-^; tg[*Ol spaca6-3xV5*]; tg[*Ol izumo1-GFP*]. Background bands are indicated with asterisks. **D** To stabilize medaka Izumo1 in zebrafish sperm, a chimeric medaka-zebrafish Izumo1 was constructed where the 4HB of zebrafish Izumo1 was replaced with the homologous medaka sequence. The model of the AlphaFold2-predicted complex as tested in the genetic experiment is shown with the medaka sequences in color and the zebrafish sequences in shades of grey (ectodomains of the relaxed top-scoring model; left). For simplicity, the complex is only shown with medaka Bouncer but is similarly predicted to form also with zebrafish Bouncer. Scoring of male fertility as in (A and B) with wild-type females in black and medaka Bouncer-presenting females in blue (right).**[1]** Data replotted from Deneke et al., 2024. **[2]** Data replotted from Binner et al., 2022. **[3]** Experiment performed with *bncr*^+/-^; tg[*mCherry-Ol bncr*] females.

Because medaka sperm is highly selective towards medaka Bouncer (Gert et al., 2021, 2023), the unexpected compatibility of medaka Spaca6 with the zebrafish fertilization machinery suggests that selectivity may be mediated by medaka Izumo1. In line with this hypothesis, medaka Izumo1 does not rescue zebrafish *izumo1* KO even in conjunction with medaka Spaca6 (**Fig. 2A**). To test whether expression of medaka Izumo1 in zebrafish sperm is sufficient for zebrafish sperm to recognize medaka Bouncer, we crossed zebrafish tg[*Ol izumo1-GFP*] males to zebrafish females presenting medaka Bouncer on their eggs. These eggs are only rarely fertilized with zebrafish sperm but can be fertilized by medaka sperm *in vitro* (∼20%) (Gert et al., 2023). However, zebrafish males expressing medaka Izumo1 did not fertilize medaka Bouncer-presenting eggs independent of the mutant background and the presence/absence of medaka Spaca6 (**Fig. 2B**).

Taken together, these results demonstrate that medaka Spaca6 – unlike medaka Izumo1 – is fully compatible in the zebrafish context both with the zebrafish sperm factors and zebrafish Bouncer on the egg.

### The Bouncer binding site of medaka Izumo1 is compatible with zebrafish Bouncer

The unexpected behavior of medaka Spaca6 and Izumo1 prompted us to investigate the expression levels of the transgenic proteins. Western blot analysis confirmed robust expression of medaka Spaca6-3xV5 in mature sperm (**Fig. 2C**). When comparing sperm to total testis samples, it appears that medaka Spaca6 is strongly enriched in mature sperm (comparing lanes 2 with 4 and 5). In contrast, medaka Izumo1-GFP is well expressed in testis but only weakly detected in mature sperm (comparing lanes 2 and 5). This suggests that medaka Izumo1 is not egiciently incorporated into the zebrafish complex in mature sperm, which could explain the observed sterility of *izumo1*^-/-^; tg[*Ol izumo1-GFP*] males.

To stabilize medaka Izumo1 in zebrafish sperm, we reasoned that only the N-terminal 4HB of Izumo1 interacts with Bouncer and as such only the 4HB of medaka Izumo1 is required to recognize medaka Bouncer. All other domains of Izumo1 could be kept as the zebrafish sequence to stabilize the complex with Spaca6 and Tmem81. We constructed such a chimeric Izumo1 and introduced it into the *izumo1*^-/-^ background as a transgene (**Fig. 2D**). The Izumo1 chimera yielded a surprising result: It partially rescued *izumo1* KO with wild-type eggs (**Fig. 2D**). While this result showed that we had indeed achieved the stabilization of the chimeric Izumo1 protein, providing the Bouncer binding site of medaka Izumo1 was unexpectedly able to bind zebrafish Bouncer when paired with zebrafish Spaca6. Mirroring the behavior of medaka Spaca6, only providing part of the medaka Bouncer binding site through the chimeric Izumo1 was not sufficient to recognize medaka Bouncer (**Fig. 2B** and **D**). This did not change even in the presence of medaka Spaca6, which would be able to complement the full medaka Bouncer binding site (**Fig. 2D**). However, in the latter scenario endogenous zebrafish Spaca6 may still be present and stabilized by the Izumo1 chimera. Therefore, we introduced the *Ol spaca6-3xV5* and *izumo1*^*chimera*^*-3xFLAG-GFP* transgenes into the *spaca6*^-/-^; *izumo1*^-/-^ double KO background to guarantee that all Spaca6 and Izumo1 in the sperm carries the medaka Bouncer binding site. However, these double transgenic, double KO males fertilized neither wild-type nor medaka Bouncer-presenting eggs (< 3%) (**Fig. 2D**). This suggests that the transgenic chimeric Izumo1 and medaka Spaca6 proteins can stabilize each other very poorly even though individually they are compatible with the zebrafish machinery.

Although we were effectively unable to provide the complete medaka Bouncer binding site on zebrafish sperm, these results show that both binding sites of medaka Bouncer on Spaca6 and Izumo1 can individually recognize zebrafish Bouncer when paired with their zebrafish counterpart.

## Discussion

### The proteins of the conserved fertilization complex also function in medaka sperm

Bouncer was discovered as an essential mediator of sperm-egg binding on zebrafish eggs, and providing its medaka ortholog on zebrafish eggs is sufficient to allow entry of medaka sperm (Herberg et al., 2018). With the recent discovery that a conserved complex of Spaca6, Izumo1 and Tmem81 on vertebrate sperm is the Bouncer receptor in zebrafish (Deneke et al., 2024), we hypothesized that this complex explains the crossing of the species barrier from the side of the sperm. To test this, we first established that medaka sperm relies on the same molecular principles of sperm-egg interaction as zebrafish. Indeed, AlphaFold2-Multimer predicted the Spaca6-Izumo1-Tmem81-Bouncer complex for the medaka orthologs, and we found that *spaca6* and *izumo1* are essential for male fertility also in medaka (**Fig. 1A** and **B**). Interestingly, co-regulation of Spaca6-Izumo1-Tmem81 in medaka sperm was not as stringent as the co-depletion in zebrafish sperm (**Fig. 1C**; Deneke et al., 2024). This finding – while subtle – suggests the individual necessity of medaka Spaca6 and Izumo1 at the time of sperm-egg interaction, whereas in zebrafish it cannot be disentangled from the necessity to stabilize the other complex members. Our sperm factor mutant analyses in medaka underscore the central role of Spaca6 and Izumo1 for fertilization not only in mammals but across broader branches of vertebrates.

### Assembling and providing Spaca6-Izumo1-Tmem81 in sperm – a delicate system

Having established that all three factors that directly interact and bridge sperm (Spaca6 and Izumo1) and egg (Bouncer) are essential for medaka fertilization (**Fig. 1**; Gert et al., 2023), we addressed the interesting question of the species compatibility with a focus on the recently uncovered Bouncer receptor: Does the substitution of Spaca6 and Izumo1 on medaka and zebrafish sperm with their orthologs enable the recognition of the heterospecific Bouncer? This poses a seemingly simple genetic experiment of ablating the endogenous genes and providing the orthologous sequences transgenically. However, we faced significant challenges in ensuring that not only the genes but also the encoded proteins were provided in the mature sperm. Currently, we have no evidence that zebrafish Spaca6 and Izumo1 are expressed at sufficient levels in medaka sperm (**Fig. 1**). We encountered a similar problem with medaka Izumo1 destabilization in zebrafish sperm (**Fig. 2C**). We propose two possible reasons for this hurdle. (1) All proteins in the sperm are produced during spermatogenesis as mature sperm is largely transcriptionally silent (Steger, 1999). Moreover, spermatogenesis is a highly specialized and strictly controlled process with major cellular rearrangements from a spheroid germ cell to a flagellated spermatozoon, which sheds most of its cellular components and organelles that are not strictly necessary for fertilization (Guraya, 1987; Schulz et al., 2010). If heterospecific Spaca6 and Izumo1 proteins were not processed and recognized as such, they would likely face the same fate as many other proteins that are not packaged into the mature sperm. (2) Protein stabilization requires the complex formation of Spaca6-Izumo1-Tmem81. Our observations from zebrafish suggest that the successful packaging of the three proteins is dependent on complex formation as single gene KO testis produces the other two proteins, which are then depleted in mature sperm (Deneke et al., 2024). Therefore, heterospecific Spaca6 and Izumo1 are likely only stable if they are integrated into the endogenous protein complex. This appears to be the case for medaka Spaca6 in the zebrafish context (**Fig. 2A** and **C**), which could be explained by Spaca6 being the most similar complex member between zebrafish and medaka (**Fig. 1A**). In contrast, the other proteins may require additional stabilization, which could be provided by additionally expressing the heterospecific Tmem81 to complete the whole trimeric complex. Overall, these observations suggest that the fertilization factors are part of a complex protein network that requires compatible factors to be presented on the mature sperm.

### Medaka Spaca6 and Izumo1 have the potential to recognize zebrafish Bouncer

In contrast to the zebrafish factors in the medaka context, we have direct evidence that medaka factors can be expressed and function in zebrafish sperm (**Fig. 2**). Surprisingly, they behave in the opposite way as initially expected. Based on the detailed analysis of Bouncer’s molecular signatures in Gert et al., 2023, we had expected medaka Spaca6 and Izumo1 to be more selective than zebrafish Spaca6 and Izumo1 as zebrafish sperm was more tolerant towards altered Bouncer sequences. However, here we find that medaka Spaca6 is fully compatible with an otherwise zebrafish sperm complex and zebrafish Bouncer without showing any preference for medaka Bouncer in this context (**Fig. 2A-C**). The same is observed for an Izumo1 variant providing the medaka Bouncer binding site (**Fig. 2D**). Unfortunately, we were not successful in conclusively supplying the complete medaka Bouncer binding site on zebrafish sperm (**Fig. 2D**). Hence, it remains an open question if such an altered sperm would prefer medaka Bouncer. Nevertheless, these experiments give an interesting indication: In fish, sperm-egg interaction through Spaca6-Izumo1-Bouncer appears to be less constrained than the cis-interactions within the sperm complex.

### The molecularly constrained sperm versus the liberated egg?

The relative tolerance to sequence divergence in fish sperm-egg recognition is consistent with the behavior of Bouncer, which incidentally is the most divergent between zebrafish and medaka among the fertilization factors (**Fig. 1A**). While Bouncer has specific molecular signatures that are required for the recognition by either zebrafish or medaka sperm, Bouncer proteins from different fish species preserve a surprisingly wide sperm compatibility across the teleost lineage despite considerable sequence divergence (Gert et al., 2023). As convincingly discussed in Gert et al., 2023, this may be due to other strong species barriers acting before or after sperm-egg interaction (e.g. habitat, mate choice, genome incompatibility) or due to the molecular promiscuity being selectively neutral or even advantageous as hybridization is common in fish, especially in cyprinids (Scribner et al., 2000; Umezawa et al., 2020).

Hence, a picture of evolutionarily constrained sperm proteins and a more flexible egg protein emerges. Considering the wider vertebrate lineage, this dichotomy can be expanded as mammalian IZUMO1 interacts with JUNO on the egg (Bianchi et al., 2014). Here, not only the protein sequence but the overall protein identity diverged between fish and mammals. How can the opposite sides of the same protein-protein interaction display such difference? We speculate that the egg ligand is more flexible because it has only one main requirement: the recognition of the sperm complex with sufficient affinity. In contrast, the sperm receptor by nature of being a protein complex must not only maintain affinity to the egg ligand but also to its interaction partners on the sperm. Given that we still lack a comprehensive picture if and how the Spaca6-Izumo1-Tmem81 complex cooperates with the other known fertilization factors, there might be additional constraints that are yet to be discovered. Consequently, the elucidation of the machinery on sperm and egg required to achieve sperm-egg fusion is bound to also provide intriguing insights into evolution of sexual selection in vertebrates.

## Acknowledgments

We thank Krista Gert for establishment and teaching of medaka husbandry and continued discussions. Proteomics analyses were performed by the Proteomics Facility at IMP/IMBA/GMI using the VBCF instrument pool, in particular Gabriela Krssakova, Karel Stejskal. We thank Krista Gert, Karin Panser, Carina Pribitzer, Theresa Humer, Victoria Deneke and Kata Szabó for technical support; Juraj Ahel and Philipp Peters for setting up an in-house AlphaFold pipeline; Anna Bandura, Gisela Deneke and Olivia Füssl for their help with genotyping; the fish facility personnel for taking care of the fish; and Victoria Deneke and the entire Pauli lab for helpful discussions about the project.

A.B. was supported by a Boehringer Ingelheim Fonds PhD fellowship. Work in the Pauli lab was supported by the FWF START program (Y 1031-B28), the ERC CoG 101044495/GaMe, an HFSP Young Investigator Award (RGY0079/2020), and the FWF SFB RNA-Deco (project number F80). The IMP receives institutional funding from Boehringer Ingelheim and the Austrian Life Sciences Program 2023 (# 48924910). For the purpose of Open Access, the authors have applied a CC BY public copyright license to any Author Accepted Manuscript (AAM) version arising from this submission.

## Author contributions

A.B., and A.P. conceived the study. A.B. designed, performed, and analyzed experiments.

A.P. supervised the study. A.B. drafted and A.B. and A.P. edited the manuscript.

## Methods

### AlphaFold2-Multimer predictions

All presented AlphaFold2-Multimer predictions were performed with the script released in Abramson et al., 2024, on the in-house computing cluster. The input sequences are the mature protein sequence of the accessions listed below. The top-scoring relaxed model and its corresponding predicted aligned error (PAE) plot was visualized with ChimeraX (Pettersen et al., 2021).

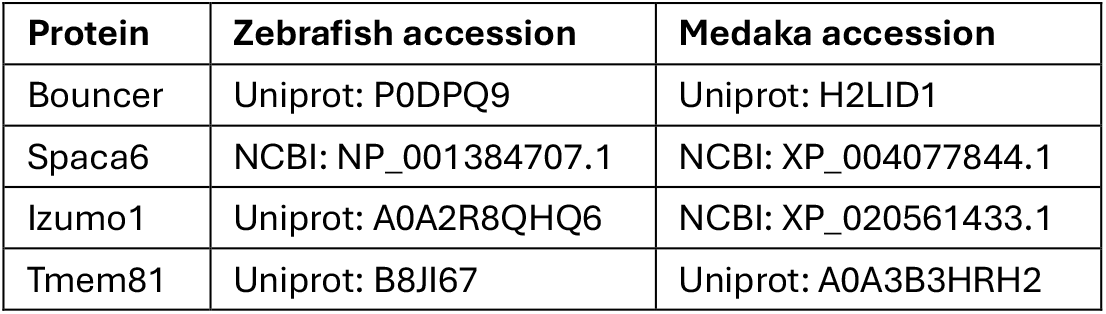

### Protein sequence alignments

The protein sequence identity and similarity were determined by sequence alignment performed with SnapGene 7.2 (GSL Biotech): Needleman-Wunch v2.4.2, BLOSUM62 matrix, Gap opening penalty 10, Gap extension penalty 1, Match score 1, Mismatch score 0. The inputs were the full-length sequences of the accessions listed above.

### Medaka and zebrafish husbandry

Medaka (*Oryzias latipes*) and zebrafish (*Danio rerio)* were raised according to standard protocols (28 °C water temperature; 14/10 h light/dark cycle). The CAB medaka strain were considered wild type. The first hybrid generation (TLAB) of AB zebrafish and the natural variant TL (Tüpfel long fin) served as wild-type zebrafish. Zebrafish embryos were cultured in E3 medium (5 mM NaCl, 0.17 mM KCl, 0.33 mM CaCl_2_, 0.33 mM MgSO_4_,10^−5^% Methylene Blue) at 28 ºC until 5 dpf. Medaka clutches were individualized, and embryos were cultured in Yamamoto’s Ringer’s solution (170 mM NaCl, 4 mM KCl, 5 mM MgCl_2_, 3 mM CaCl_2_, adjusted to pH 7.3 with NaHCO_3_) at 28 ºC until hatching. All fish experiments were conducted according to Austrian and European guidelines for animal research and approved by the Amt der Wiener Landesregierung, Magistratsabteilung 58 -Wasserrecht (protocols GZ 342445/2016/12, MA 58-221180-2021-16 and GZ 198603/2018/14). The full list of fish lines utilized in this study are listed in **Table 1**.

**Table 1.** Fish lines utilized in this study.

| **Medaka fish line** | **Source** |
| --- | --- |
| *spaca6*<sup>-/-</sup> | This study |
| tg[*dcst2:Dr spaca6-3xV5; Dr cmlc2:EGFP*] | This study |
| *izumo1*<sup>-/-</sup> | This study |
| tg[*dcst2:Dr izumo1-3xFLAG-sfGFP; Dr cmlc2:EGFP*] | This study |
| *bncra*<sup>-/-</sup> | Gert et al., 2023 |
| tg[*ubi:sfGFP-Dr bncr*] | Gert et al., 2021 |
| **Zebrafish fish line** | **Source** |
| *bncr*<sup>-/-</sup> (vbc1; ZFIN:ZDB-ALT-181207-1) | Herberg et al., 2018 |
| tg[*actb2:sfGFP-bncr(Medaka Top)*] | Gert et al., 2023 |
| tg[*ubi:mCherry-Ol bncr*] | This study |
| *spaca6*<sup>-/-</sup> (vbc12; ZFIN:ZDB-ALT-220110-7) | Binner et al., 2022 |
| tg[dcst2:Ol spaca6; cres:EGFP] | This study |
| tg[dcst2:Ol spaca6-3xV5; cres:EGFP] | This study |
| izumo1<sup>-/-</sup> | Deneke et al., 2024 |
| tg[dcst2:Ol izumo1-sfGFP; cmlc2:EGFP] | This study |
| tg[dcst2:Ol izumo1[aa1-119]-Dr izumo1[127-336]-3xFLAG-sfGFP; cmlc2:EGFP] | This study |

### Generation of testis cDNA

To sequence and clone *spaca6* and *izumo1* cDNA, the testes of one medaka were dissected. Total RNA was isolated with Trizol (Ambion) and reverse-transcribed using the iScript cDNA Synthesis Kit (BioRad).

### Generation of *spaca6* and *izumo1* knock-out medaka

Knock-out medaka were generated using CRISPR/Cas9-mediated mutagenesis. Guide RNAs (sgRNAs, listed below) were synthesized according to Gagnon et al., 2014, using the MEGAscript T7 Transcription Kit (Thermo Fisher). Two sgRNAs and Cas9 protein (in-house) were co-injected into one-cell CAB embryos according to Rembold et al., 2006, and raised until adult. The adult fish (F_0_) were crossed to wild-type fish. The offspring (F_1_) were genotyped by PCR (primers listed below) on gDNA isolated from fin clips. InDel (insertion-deletion) alleles were identified by agarose gel electrophoresis. Heterozygous fish (F_1_) with the same mutant allele were incrossed. Homozygous offspring (F_2_) were genotyped to determine the nature of the mutation by Sanger sequencing (**Fig. S1**). To identify the mutant mRNA, *izumo1* and *spaca6* cDNA was amplified from homozygous mutant testes cDNA by PCR (primers listed below) and characterized by Sanger Sequencing (**Fig. S1**).

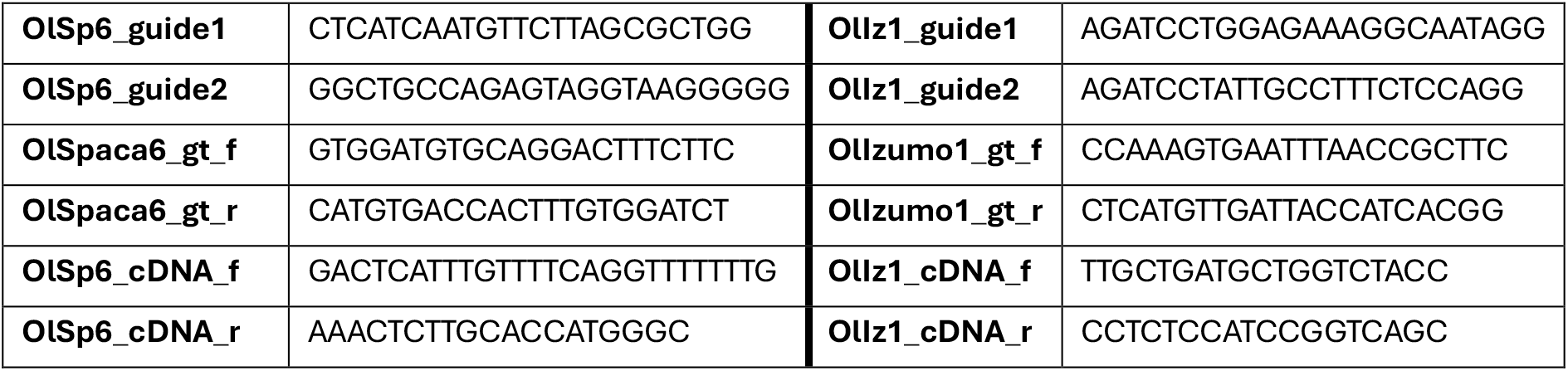

### Cloning of transgenes introduced into medaka

Transgenes were cloned by Gibson assembly. The upstream sequence of *Ol dcst2* (ASM223467v1, chr16:31790185-31790869) was amplified by PCR from gDNA as putative medaka promoter sequence driving testis expression in analogy of the zebrafish *dcst2* promoter sequence having been shown to drive testis expression in zebrafish (GRCz11, chr16:28726943-28727663) (Deneke et al., 2024). To generate *Dr spaca6-3xV5*, the *Dr spaca6* coding sequence (CDS) was amplified from the transgene generated in Binner et al., 2022, introducing *3xV5* with primers. *Dr izumo1-3xFLAG-sfGFP* was amplified from the transgene generated in Deneke et al., 2024. To construct *Ol dcst2:Dr spaca6-3xV5* and *Ol dcst2:Drizumo1-3xFLAG-sfGFP*, the promoter fragment and the respective coding fragment were introduced into a vector with a *Dr cmlc2:eGFP* reporter flanked by I-SceI meganuclease sites.

### Generation of transgenic medaka lines

Transgenic medaka were generated by meganuclease-mediated transgenesis according to Rembold et al., 2006. Briefly, the transgene-containing plasmid was digested with I-SceI for 1 h at room temperature. The restriction reaction was injected into one-cell CAB embryos. Once adult, the fish were crossed to wild-type fish and the offspring was assessed for fluorescent hearts. Each fish line was expanded from a single founder. To obtain double transgenic fish, tg[*Ol dcst2:Dr spaca6-3xV5*] and tg[*Ol dcst2:Drizumo1-3xFLAG-sfGFP*] fish were crossed and screened for strong heart fluorescence. Juvenile fish were genotyped for the presence of both transgenes by fin clips and a PCR with transgene-specific primers (listed below).

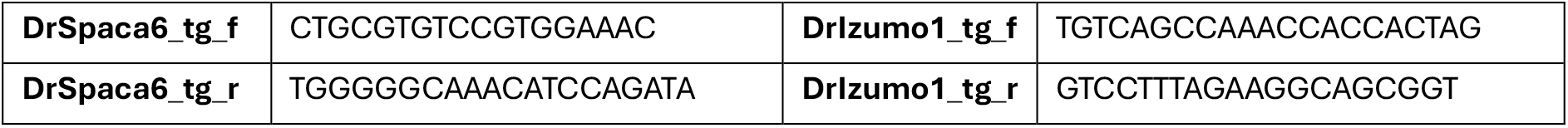

### Assessment of male medaka fertility

To quantify male fertility in medaka, one male was co-housed with two to four females. Once a day, all eggs were collected. In the afternoon, embryos with cellular cleavage were considered fertilized. The ratio of fertilized embryos to total number of eggs collected from one breeding tank during the observation period is reported as the fertilization rate per male.

### Medaka and zebrafish sperm collection

To collect unactivated fish sperm, one zebrafish or medaka male was anesthetized using 0.01% tricaine and placed in a wet sponge under a dissecting microscope. By applying gentle pressure, sperm was expelled and collected into a glass capillary by mouth pipetting. The sperm was stored on ice in Hank’s saline solution (5.4 mM KCl, 0.137 M NaCl, 1 mM MgSO_4_, 4.2 mM NaHCO_3_, 0.25 mM Na_2_HPO_4_, 1.3 mM CaCl_2_).

### Relative protein quantification by targeted mass spectrometry via parallel reaction monitoring

Relative protein abundance was determined as described in Deneke et al., 2024. Briefly, 2.5 - 3.5 × 10^7^ medaka sperm per biological replicate were collected and lysed in SDT buger (4% SDS, 100 mM DTT, 100 mM Tris-HCl pH 7.5, 1 mM MgCl_2_ + benzonase (Merck)) for 1 h at room temperature, then boiled for 10 min at 95 ºC before sonication. The protein extract was mixed with 200 ml of 8 M urea in 0.1 M TEAB pH 8.5 and digested with trypsin (1 part protease per 20 parts protein; w/w) using the FASP method. The final peptide concentration was determined by separating an aliquot on a LC-UV system. Nano-LC-MS/MS analysis was performed on a nano-HPLC system (Vanquish Neo UHPLC-System). The Orbitrap Exploris 480 mass spectrometer was operated by a mixed MS method, which consisted of one full scan followed by the PRM of targeted peptides. A scheduled PRM method (sPRM) development, data processing and manual evaluation of results were performed in Skyline. Spectra of unique peptides of the proteins of interest (medaka Izumo1, Spaca6, Tmem81) and the ‘control proteins’ (medaka proteins considered constant) were recorded. All peptides included in the final sPRM method are listed below.

The sum of the peptide areas of each normalization protein was used to obtain a normalization factor per sample. After normalization to the control proteins, the sum of peptide areas of each protein of interest was averaged and normalized to the mean of the wild-type samples.

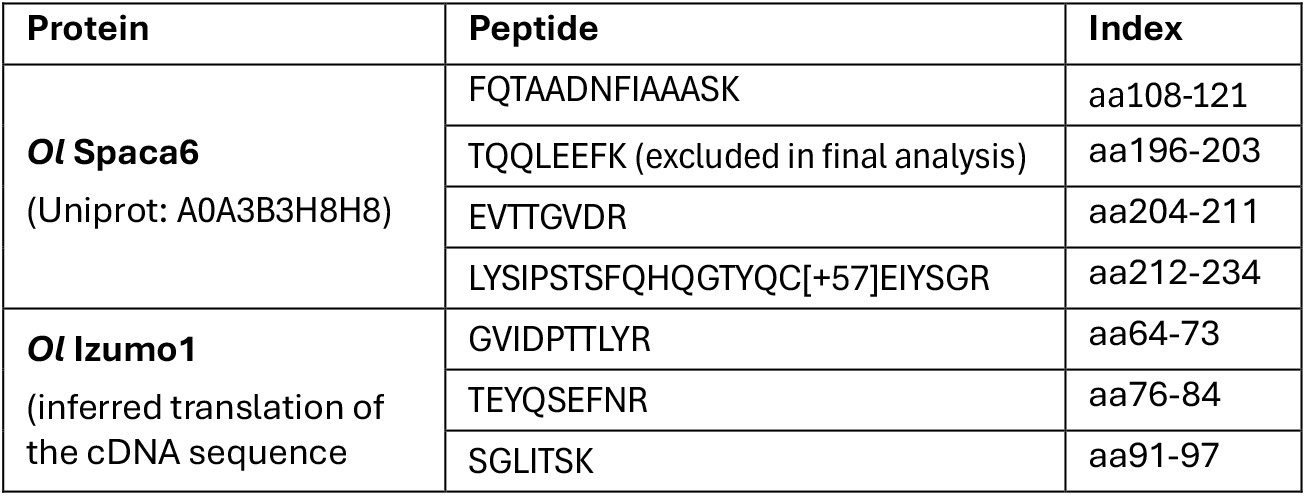
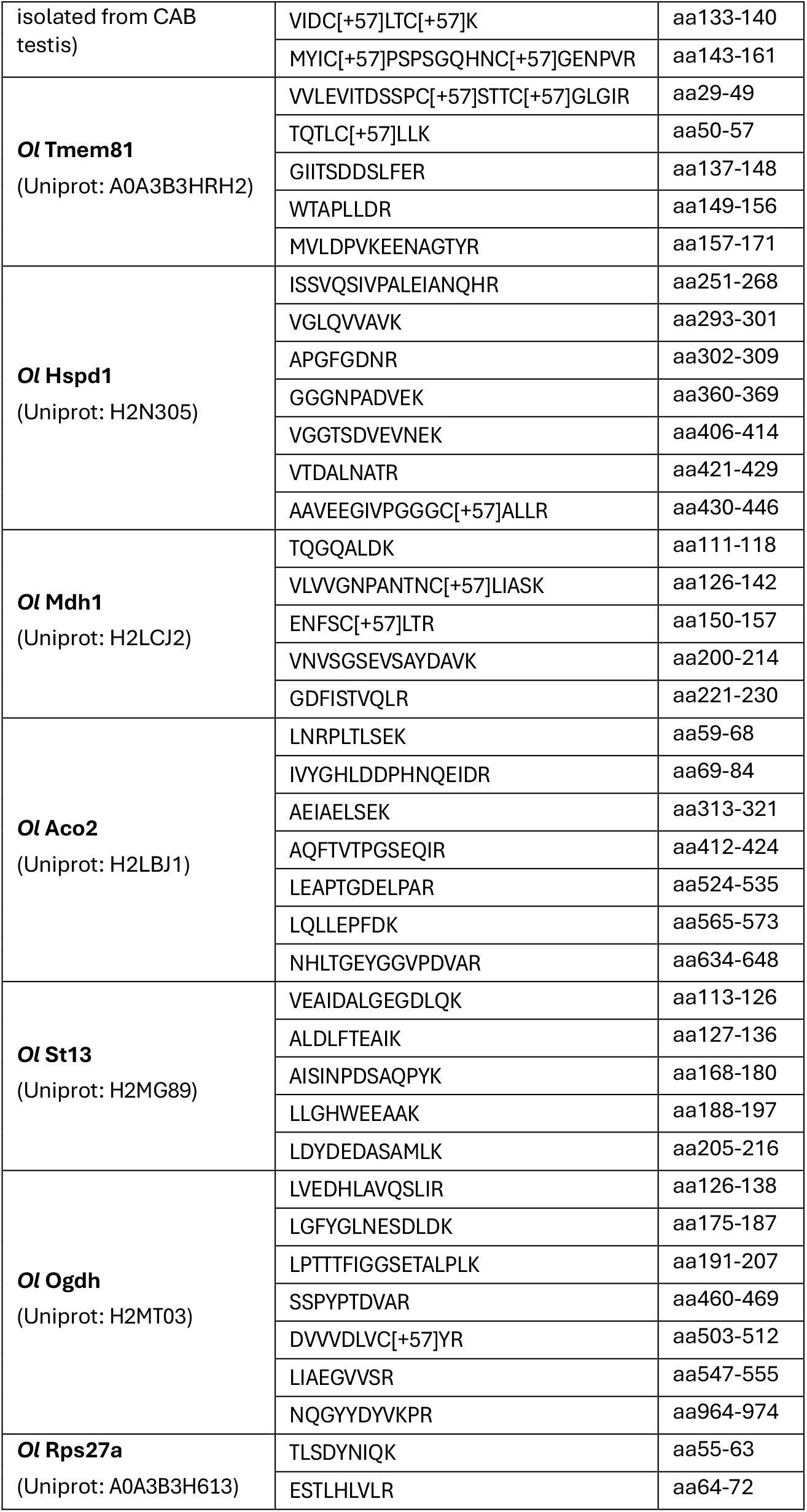

### *In vitro* fertilization with zebrafish eggs

Comparative *in vitro* fertilization was performed as in Gert et al., 2023. Briefly, zebrafish eggs were collected by anesthetizing a female with 0.1% tricaine and expelling 50 - 100 eggs into a dry petri dish by gentle pressure on the belly. This was repeated in a second petri dish with the same female. Immediately, 45 µL of zebrafish or medaka sperm (50,000 - 100,000 µL^-1^) was added to each dish and the gametes were activated with 500 µL E3 medium. After 5 min, the embryos were cultured as usual.

### Cloning of transgenes introduced into zebrafish

Transgenes were cloned by Gibson assembly. The *Ol spaca6* and *Ol izumo1* CDS’s were amplified from testis cDNA. The *Dr dcst2:Ol spaca6* transgene was constructed by introducing *Ol spaca6* CDS into a vector generated in Deneke et al., 2024, containing the *Dr dcst2* promoter and a *cres:eGFP* reporter flanked by Tol2 recognition sites. *Dr dcst2:Ol spaca6-3xV5* was constructed the same way but with primers encoding 3xV5. Similarly, *Dr dcst2:Ol izumo1-sfGFP* was constructed with an *Ol izumo1* fragment and a fragment encoding sfGFP into the Tol2 vector that contains a *cmlc2:eGFP*. The *izumo1*^*chimera*^ transgene was generated by subcloning the *Ol izumo1* and *Dr izumo1* transgenes into the same vector with primers that assemble *Dr dcst2:Ol izumo1*[aa1-119]*-Dr izumo1*[127-336]*-3xFLAG-sfGFP*.

### Generation of transgenic zebrafish

Transgenic zebrafish were generated analogous to Deneke et al., 2024. Briefly, *tol2* mRNA was co-injected with the transgene-containing vector into one-cell zebrafish embryos. *Spaca6* transgenes were injected into *spaca6*^+/-^ embryos and *izumo1* transgenes into *izumo1*^+/-^ embryos. To identify founder fish, *tol2*-injected adult males were crossed to *spaca6*^-/-^ or *izumo1*^-/-^ females. Offspring with fluorescent neural crest (tg[*spaca6*]) or fluorescent hearts (tg[*izumo1*]) were raised. All transgenic lines were expanded from a single founder. To obtain double transgenic fish (tg[*spaca6*]; tg[*izumo1*]), the transgenic lines were crossed and progeny with fluorescent neural crest and hearts were raised.

### Assessment of male zebrafish fertility

Each assessed male was crossed with two to three females. At ∼5 hpf, the number of developing and non-developing embryos was counted. The fraction of embryos with cell cleavages (considered fertilized) is the fertilization rate of the clutch.

### Western blot

Testes of one male were dissected and lysed in 150 µL SDT-PAGE buffer (4% SDS, 100 mM DTT, 100 mM Tris-HCl pH 7.5, 1 mM MgCl_2_, 10% glycerol + bromophenol blue + benzonase (Merck)) and homogenized with a plastic pestle before incubation for 1 h at room temperature. 2.5 × 10^7^ sperm were lysed in 35 µL SDT-PAGE for 1 h at room temperature. All protein samples were boiled for 10 min at 95 ºC. SDS-PAGE was performed with precast gels (Any kD Mini-PROTEAN TGX, BioRad) and blotted onto nitrocellulose membranes by wet transfer. The blot was blocked with 2.5% milk powder in TBST (Tris-buffered saline + 0.1% Tween 20) before incubation with primary antibody (listed below) in blocking buffer overnight at 4 ºC. After washing with TBST, the blot was incubated with HRP-conjugated secondary antibody (listed below) in blocking buffer for 45-60 min at room temperature. After washing, the blot was developed with Clarity Western ECL Substrate (BioRad) in a ChemiDoc (BioRad). The GFP blot was stripped (Restore Western Blot Stripping Buffer, ThermoFisher) and reprobed for eEF2 as a loading control.

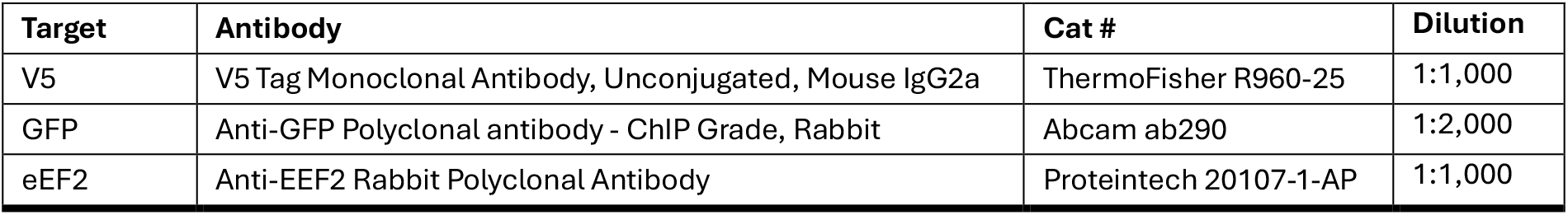
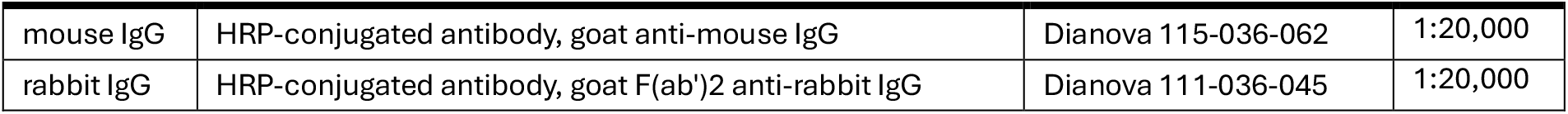

| Target | Antibody | Cat # | Dilution |
| --- | --- | --- | --- |
| V5 | V5 Tag Monoclonal Antibody, Unconjugated, Mouse IgG2a | ThermoFisher R960-25 | 1:1,000 |
| GFP | Anti-GFP Polyclonal antibody - ChIP Grade, Rabbit | Abcam ab290 | 1:2,000 |
| eEF2 | Anti-EEF2 Rabbit Polyclonal Antibody | Proteintech 20107-1-AP | 1:1,000 |
| mouse IgG | HRP-conjugated antibody, goat anti-mouse IgG | Dianova 115-036-062 | 1:20,000 |
| rabbit IgG | HRP-conjugated antibody, goat F(ab')<sub>2</sub> anti-rabbit IgG | Dianova 111-036-045 | 1:20,000 |

## Supplemental information

**Figure S1.**
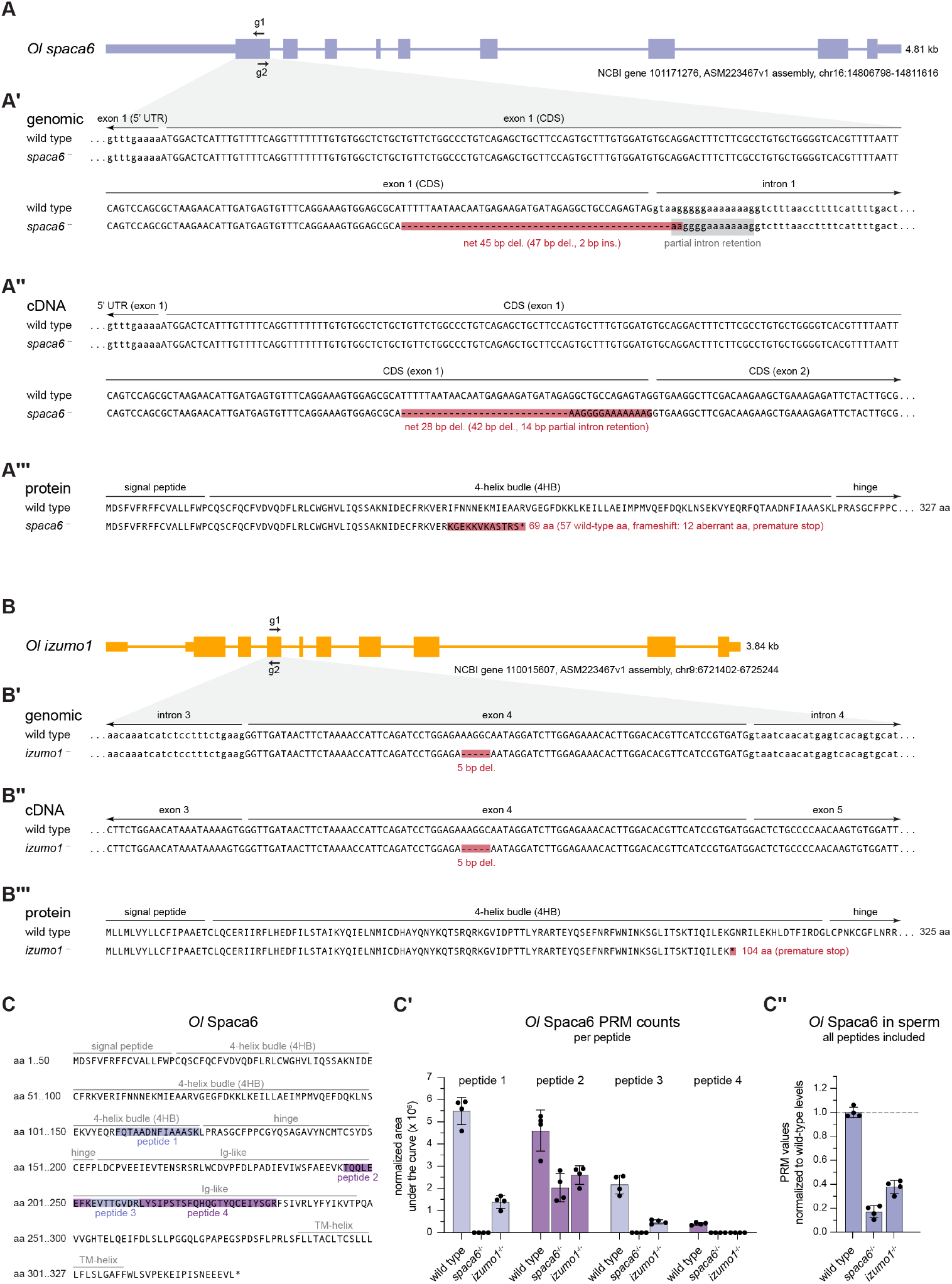
Characterization of the medaka *spaca6* and *izumo1* KO alleles. **A** Characterization of the *Ol spaca6* mutant. Schematic representation of the gene body of medaka *spaca6*; thick lines: coding exons; medium lines: UTR exons; thin lines: introns. The position of the gRNAs (g1, g2) in exon 1 are indicated. **A’** Excerpt of the exon1 highlighting the nature of the CRISPR/Cas9-induced mutation (red) and the intronic sequence (grey), that is included in the mutant mRNA. **A’’** Alignment of the sequenced mutant cDNA against the wild-type cDNA highlighting the mutated sequence and altered splicing in red. **A’’’** Translation of the mutant cDNA leading to an early frameshift and premature stop. **B** Characterization of the *Ol izumo1* mutant. A 5 bp deletion in exon 5 (**B’**) and the cDNA (**B’’**) leads to an early premature stop (**B’’’**). **C** Extended data related to Fig. 1C. Amino acid (aa) sequence of medaka Spaca6 with domain annotations in grey and the four tryptic peptides quantified by PRM in shades of purple. **C’** Raw counts of Spaca6 in medaka sperm per peptide. Comparing the trend of all peptides, peptide 2 was excluded as an unexplained outlier in the final quantification (Fig. 1C). Considering that all four peptides are encoded after the premature stop and peptides 2-4 are consecutive, our leading hypothesis is that peptide 2 is not a unique peptide although it is predicted to be unique when searching against an *in silico* tryptic digest of the HdrR reference genome. This is plausible because the tested CAB strain carries numerous SNP-based amino acid variations compared to the HdrR reference. Therefore, peptide 2 may represent a mixed signal of Spaca6 and an unidentified protein present in CAB sperm. **C’’** Quantification of Spaca6 protein analogous to Fig. 1C when including all four peptides.

